# Genome assembly of the acoel flatworm *Symsagittifera roscoffensis*, a model for research on photosymbiosis

**DOI:** 10.1101/2022.08.27.505549

**Authors:** Pedro Martinez, Kirill Ustyantsev, Mikhail Biryukov, Stijn Mouton, Liza Glasenburg, Simon G. Sprecher, Xavier Bailly, Eugene Berezikov

## Abstract

*Symsagittifera roscoffensis* is a well-known member of the order Acoela that lives in symbiosis with the algae *Tetraselmis convolutae* during its adult stage. Its natural habitat is the eastern coast of the Atlantic, where at specific locations thousands of individuals can be found lying in large pools on the surface of sand at low tide and in the sandy interstitial web at high tide. As a member of the Acoela it has been used as a proxy for early bilaterian animals; however, its phylogenetic position remains debated. In order to understand the basic structural characteristics of the acoel genome, we sequenced and assembled the genome of aposymbiotic *S. roscoffensis*. The size of *S. roscoffensis* genome was measured to be in range 910 - 940 Mb. Sequencing of the genome was performed using PacBio Hi-Fi technology. Hi-C and RNA-seq data were also generated to scaffold and annotate the genome. The resulting assembly is 1.1 Gb large (covering 118% of the estimated genome size) and highly continuous, with N50 scaffold size of 1.04 Mb. The repetitive fraction of the genome is 61%, of which 85% (half of the genome) are LTR retrotransposons. Genome-guided transcriptome assembly identified 34,493 genes, of which 29,351 are protein coding (BUSCO score 97.6%), and 30.2% of genes are spliced leader (SL) trans-spliced. The completeness of this genome suggests that it can be used extensively to characterize gene families and conduct accurate phylogenomic reconstructions.

**Significance:** *Symsagittifera* is a representative of the phylum Acoela, the first offshoot of bilaterian animals. This key phylogenetic position adds an extra value to the knowledge of its genome, since it will inform us on how the genome of a bilaterian ancestor might have looked like. Moreover, *Symsagittifera roscoffensis* is a model organism used in symbiogenesis research. Host and algae can be cultured independently and, after mixing, the symbiosis can be followed. Symbiogenesis was established early on during the evolution of Metazoa. In spite of its biological relevance, very little is known on the molecular mechanisms that control it. Here the genome of the acoel host should provide us with insights on the first adaptations to symbiogenesis occurring in bilateral animals.

## Introduction

Acoel flatworms (order Acoela) are members of the phylum Xenacoelomorpha, which also include the clades Nemertodermatida and Xenoturbellida (Hejnol et al. 2009). Today, the acoels are represented by approximately 400 described species, almost all of which are marine (Philippe et al. 2011). They exhibit a remarkable anatomical diversity, with many having salient characteristics such as an association with photosymbionts or extensive regenerative abilities. *Symsagittifera roscoffensis*, a species with the aforementioned properties, is one of the best-studied species of the Acoela (Figure 1).

**Figure 1.**
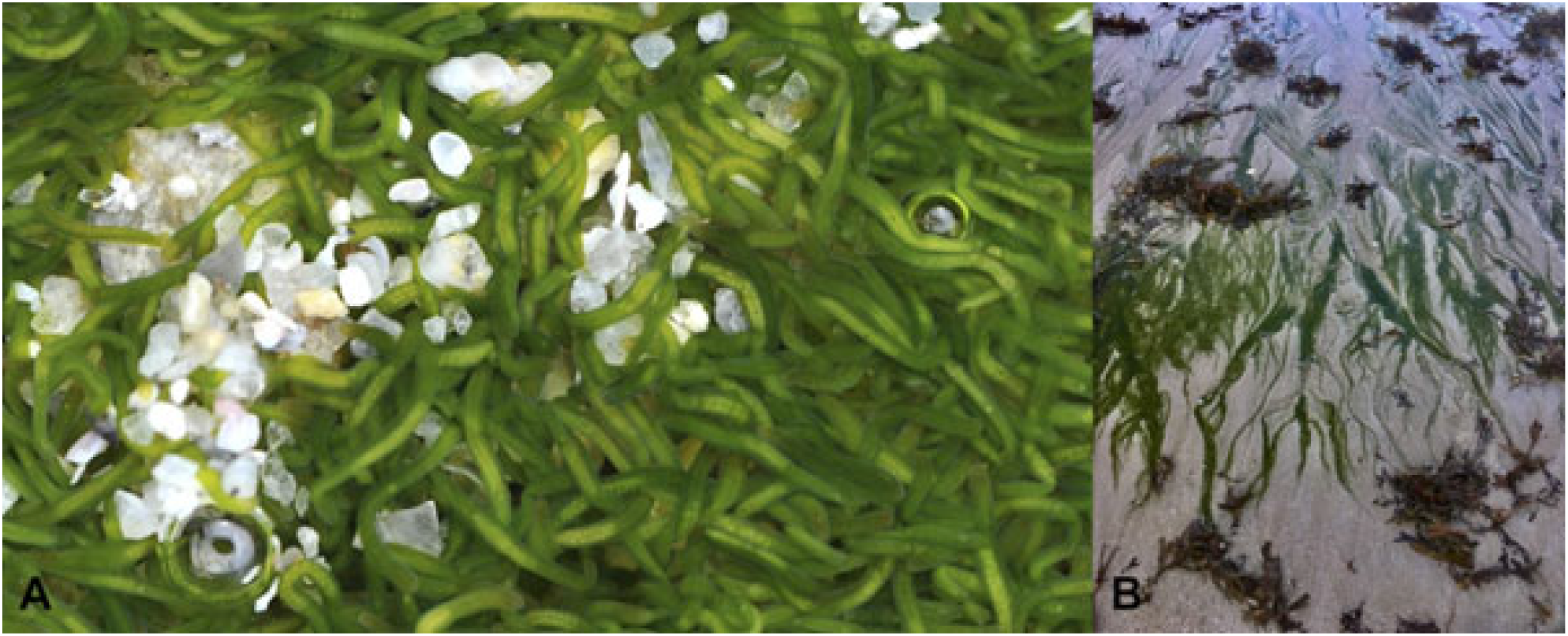
Specimens of the acoel *S. roscoffensis* in their natural habitat. **A**. Adult (gravid) *S. roscoffensis*. Credit Wilfried Thomas / Station Biologique de Roscoff. **B**. Biotope. Pools of adult specimens at low tide in a Brittany beach (France).

Photosymbiotic adult stage is abundant along most of the Atlantic coast of Europe (from Wales to Gibraltar), easy to collect, and lives in an obligatory relationship with the algae *Tetraselmis convolutae*, an association that relies on the interchange of key metabolites (Bailly et al. 2014). The algae are acquired from the environment after the worms have hatched. This event can be reproduced in the laboratory, making *S. roscoffensis* an ideal model to study the establishment and molecular control of symbiosis (Arboleda et al. 2018). These worms also show an extensive capacity to regenerate; this mostly occurs in the anterior part of the body, signifying the regeneration of the entire brain from scratch and providing a nice model for understanding the regenerative capacities of the central nervous system (Sprecher et al. 2015). These practical advantages of *S. roscoffensis* provide a general idea of the relevance of this marine worm in different areas of biology. Yet, this is not the most salient feature of this species. As a member of the Acoela, a lineage considered to be an early offshoot of the Bilateria, it has also been used to infer how the ancestral bilaterians might have looked. However, the position of Acoela as the sister group of the remaining Bilateria (Nephrozoa) has been challenged during the last few years, resulting in some uncertainty as to its phylogenetic placement (Cannon et al. 2016; Philippe et al. 2019).

The previously discussed practical and theoretical considerations of *S. roscoffensis* and Acoela suggest that understanding the genomic characteristics of this species is of special relevance, providing us with tools to understand, for instance, symbiogenesis, regenerative processes, and the phylogenetic position of this and other acoels. Obtaining a complete genome should be of the utmost importance, as it is a vital source of information on gene content and genomic architecture. Such information could potentially be used to determine the fate and biological role of gene families and provide alternative characters (synteny, indels, etc.) for phylogenomic reconstruction.

In the recent past, we and others generated the first draft genome of *Symsagittifera roscoffensis* (Philippe et al. 2019). This was used for the characterization of several gene families and to re-analyze the affinities of the acoels. The first draft genome was relatively poor, as most contigs were very small in size (N50 < 5 kb). Here we present a new draft of the *S. roscoffensis* genome, generated using PacBio Hi-Fi technology and scaffolded with Hi-C data, which allowed us significantly increase genome assembly continuity (N50 scaffold size 1 Mb). This newly assembled genome has also provided a better estimate of *S. roscoffensis* genome size, which we determined to be 1.1 Gb. We discuss these findings in the context of the few published acoel genomes, those of the species *Hofstenia miamia* and *Praesagittifera naikaiensis* (Gehrke et al. 2019; Arimoto et al. 2019).

## Results

### Genome assembly and evaluation

In order to distinguish the genomes of *S. roscoffensis* and its microsymbiotic algae, in this study we used aposymbiotic juveniles that had not yet acquired microsymbionts (from embryos grown in artificial sea water). Based on the relative fluorescence of the PI labelling from multiple measurements, the genome size of *S. roscoffensis* is estimated to be in range of 910-940 Mb (Supplementary Figure 1). We sequenced the genome to 20x coverage with Pacific Biosciences Hi Fidelity reads (1.29 mln ccs reads, mean length 15.7 kb). For the initial genome assembly, we tested several assemblers that are designed to work with HiFi reads (Table 1), including FALCON (Chin et al. 2016), Flye (Kolmogorov et al. 2019), HiCanu (Nurk et al. 2020), Hifiasm (Cheng et al. 2021), IPA (Sovic 2022), Peregrine (Chin and Asif Khalak 2019), Raven (Vaser and Šikić 2021) and wtdbg2 (Ruan and Li 2020). For the evaluation of the assemblies, we examined: the assembly size, N50 contig length, the fraction of transcripts from *de novo* transcriptome assembly mapping to the genome, and the number of BUSCO gene models identified. Since genome sequencing was performed on a population of animals obtained directly from a natural habitat, it is expected that a substantial level of heterozygosity is present in the sequencing data and thus genome assemblers that do not collapse heterozygous regions will produce assemblies normally larger than the measured genome size. Indeed, FALCON, Flye, Hifiasm, IPA and Peregrine produced redundant assemblies of 1.2 Gb - 2.5 Gb in size (Table 1). At the same time, HiCanu and Raven over-collapsed the assemblies (737 Mb and 555 Mb), while Wtdbg2 generated assembly closest in size to the measured genome size (935 Mb). Wtdbg2 also produced the highest N50 length (140.3 Kb) among the tested assemblers. However, the fraction of de novo transcripts mapped and BUSCO models identified was the highest for the Peregrine assembly (Table 1).

**Table 1.**
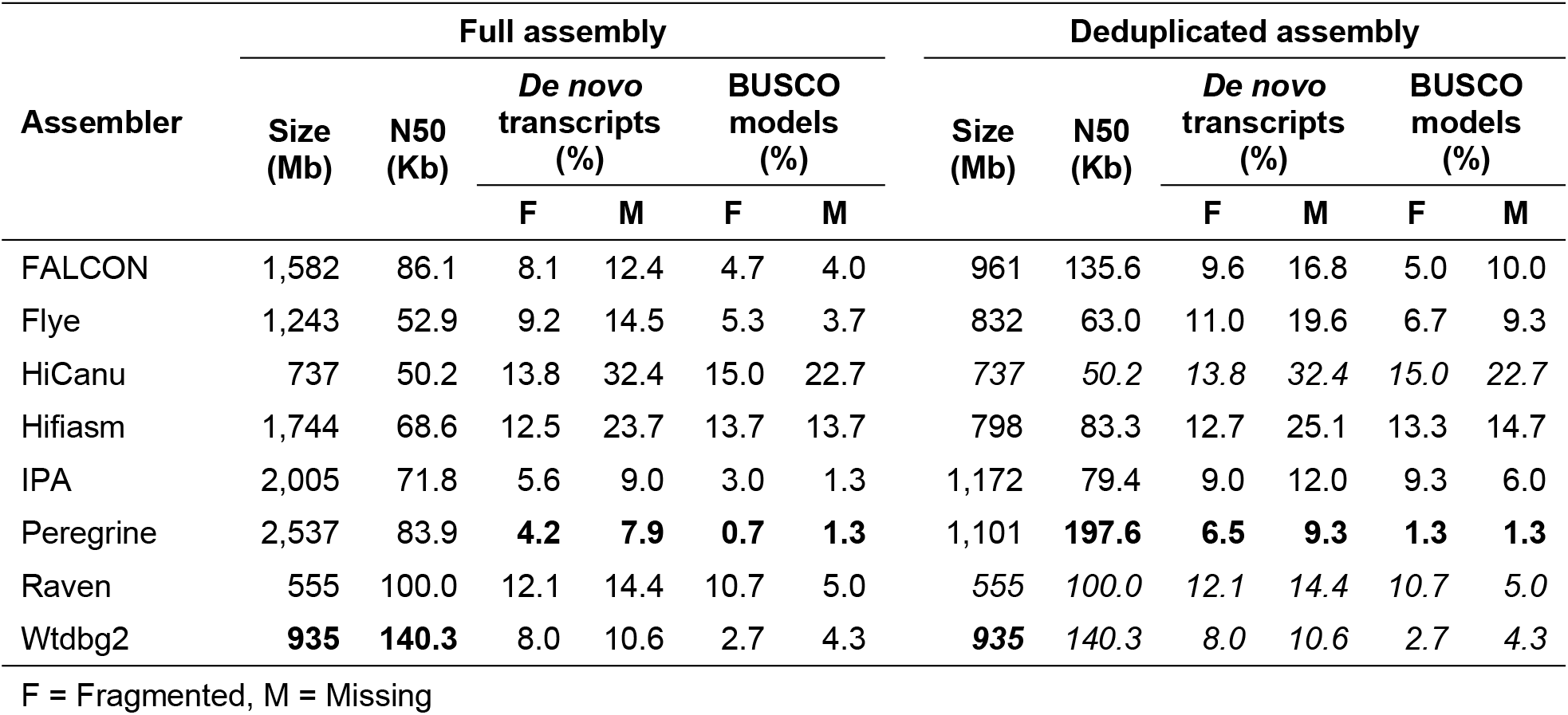
Statistics of the initial genome assemblies.

| Assembler | Full assembly |  |  |  |  |  | Deduplicated assembly |  |  |  |  |  |
| --- | --- | --- | --- | --- | --- | --- | --- | --- | --- | --- | --- | --- |
|  | Size (Mb) | N50 (Kb) | *De novo* transcripts (%) |  | BUSCO models (%) |  | Size (Mb) | N50 (Kb) | *De novo* transcripts (%) |  | BUSCO models (%) |  |
|  |  |  | F | M | F | M |  |  | F | M | F | M |
| FALCON | 1,582 | 86.1 | 8.1 | 12.4 | 4.7 | 4.0 | 961 | 135.6 | 9.6 | 16.8 | 5.0 | 10.0 |
| Flye | 1,243 | 52.9 | 9.2 | 14.5 | 5.3 | 3.7 | 832 | 63.0 | 11.0 | 19.6 | 6.7 | 9.3 |
| HiCanu | 737 | 50.2 | 13.8 | 32.4 | 15.0 | 22.7 | 737 | 50.2 | 13.8 | 32.4 | 15.0 | 22.7 |
| Hifiasm | 1,744 | 68.6 | 12.5 | 23.7 | 13.7 | 13.7 | 798 | 83.3 | 12.7 | 25.1 | 13.3 | 14.7 |
| IPA | 2,005 | 71.8 | 5.6 | 9.0 | 3.0 | 1.3 | 1,172 | 79.4 | 9.0 | 12.0 | 9.3 | 6.0 |
| Peregrine | 2,537 | 83.9 | **4.2** | **7.9** | **0.7** | **1.3** | 1,101 | **197.6** | **6.5** | **9.3** | **1.3** | **1.3** |
| Raven | 555 | 100.0 | 12.1 | 14.4 | 10.7 | 5.0 | 555 | 100.0 | 12.1 | 14.4 | 10.7 | 5.0 |
| Wtdbg2 | **935** | **140.3** | 8.0 | 10.6 | 2.7 | 4.3 | **935** | **140.3** | 8.0 | 10.6 | 2.7 | 4.3 |

F = Fragmented, M = Missing

While the *de novo* transcriptome assembly was also generated from heterozygous data and thus the higher mapping rate is expected for assemblies that do not collapse heterozygous regions, the fraction of identified BUSCO genes ideally should not be lower in the assemblies that collapse heterozygous regions. After deduplication of the redundant assemblies with purge_dups (Guan et al. 2020), the Peregrine assembly appeared to be substantially better compared to all other tested assemblies, with N50 size of 197.6 kb and the lowest fraction of missing *de novo* and BUSCO transcripts, while its assembly size of 1.1 Gb is only ~9% larger than the measured genome size (Table 1). Therefore, we decided to use the Peregrine assembly for further scaffolding and gap closing.

For genome scaffolding, we generated 388 mln Illumina read pairs (~100x genome coverage) from a Hi-C library constructed with Arima HiC+ kit. Hi-C scaffolding was performed by SALSA2 and substantially improved assembly continuity (4,475 scaffolds, N50=688.1 kb). We next scaffolded this assembly by P_RNA_scaffolder (Zhu et al. 2018) using paired-end RNA-seq reads, further improving assembly continuity (3,460 scaffolds, N50=1039.9 kb). Finally, we closed gaps in the assembly by LR_gapcloser (Xu et al. 2019) using initial PacBio Hi-Fi reads and polished the assembly with pilon (Walker et al. 2014) and RNA-seq data, reducing the number of contigs from 8,943 to 7,843 and improving N50 contig size from 197.6 kb to 237.9 kb. The parameters of the resulting genome assembly SymRos_1_5 are summarised in Table 2.

**Table 2.** Characteristics of genome assembly SymRos_1_5

|  | **Contigs** | **Scaffolds** |
| --- | --- | --- |
| Total number | 7,843 | 3,460 |
| Total length, bp | 1,101,399,379 | 1,103,025,803 |
| Average length, bp | 140,431 | 318,794 |
| Shortest, bp | 12,747 | 12,747 |
| Longest, bp | 1,836,468 | 8,003,794 |
| N50, bp | 237,875 | 1,039,899 |
| L50 | 1,417 | 287 |
| GC content, % | 36.7 | 36.7 |

### Mitochondrial genome

The mitochondrial genome of *S. roscoffensis* was previously published (Mwinyi et al. 2010). We have independently reconstructed the sequence of the mitochondrial genome from PacBio Hi-Fi genome assemblies by searching for contigs that match to mitochondrial protein sequences from another acoel flatworm, *Isodiamtera pulchra*, and also circularizing the identified contig on the internal terminal repeated sequence. The mitochondrial genome identified in this study is 99% identical to the published sequence, thus providing a good control for the quality of our nuclear genome sequencing and assembly.

### Repeat annotation

The genome is highly repetitive, with various transposable elements and simple repeats comprising more than 60% of the total genome size (Table 3). LTR retrotransposons account for 85% of all repeat elements and comprise 51% of the genome. Major prevalence of LTR retrotransposons was also previously shown for two other acoel genomes, *Hofstenia miamia* (total repeat content 53%) (Gehrke et al. 2019) and *Praesagittifera naikaiensis* (total repeat content 70%) (Arimoto et al. 2019).

**Table 3.**
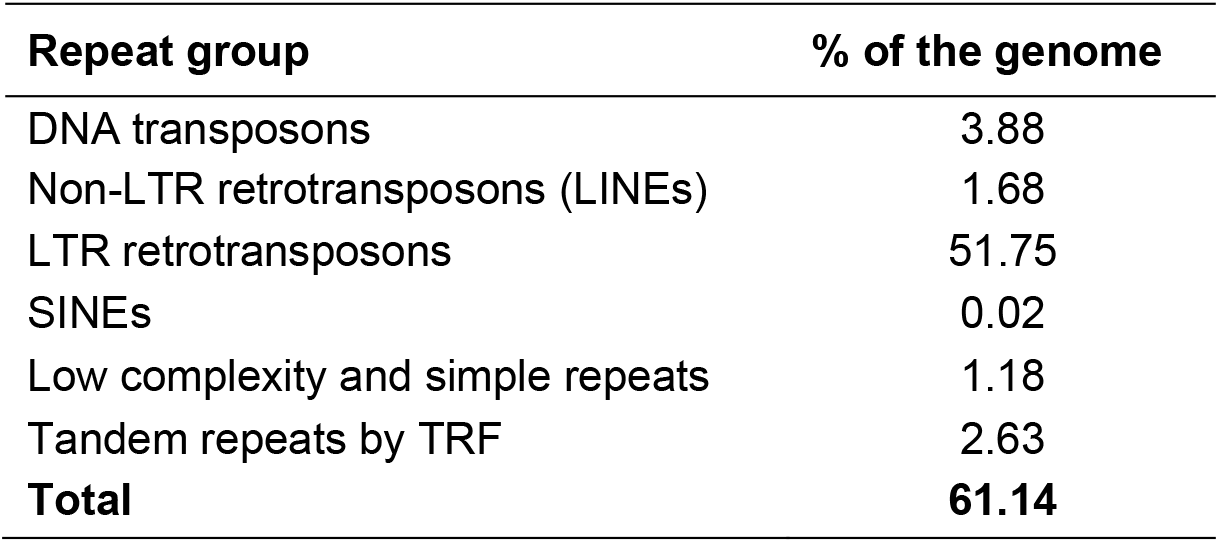
Repeat content of SymRos_1_5 genome assembly

| **Repeat group** | **% of the genome** |
| --- | --- |
| DNA transposons | 3.88 |
| Non-LTR retrotransposons (LINEs) | 1.68 |
| LTR retrotransposons | 51.75 |
| SINEs | 0.02 |
| Low complexity and simple repeats | 1.18 |
| Tandem repeats by TRF | 2.63 |
| **Total** | **61.14** |

### Gene annotation

To annotate genes in the assembly, we used TBONE (Transcript Boundaries based ON experimental Evidence) pipeline (Wudarski et al. 2017), public paired-end RNA-seq data (SRR5760179, 61.8 mln Illumina read pairs; SRR8506641, 37.4 mln Illumina read pairs) and also generated additional RNA-seq dataset from juveniles (11.9 mln Illumina reads) using Smart-3SEQ approach (Foley et al. 2019), in which 3’ ends of transcripts are sequenced. TBONE pipeline takes into account potential effects of spliced leader (SL) trans-splicing, which is present in flatworms (Ustyantsev and Berezikov 2021). By mapping *de novo* transcriptome assembly SYMROS200831 to the genome assembly and analysing non-mapped 5’-end sequences, we determined that SL trans-splicing is also present in *S. roscoffensis* and that the predominant spliced leader sequence is GCCTAATTGTTGTGATAAACTTATTAAATAGA, originating from multiple SL RNA genes in the genome assembly (Supplementary Fig. 2). Mapping RNA-seq reads that contain this leader sequence to the genome assembly allowed identification of transcripts that undergo SL trans-splicing and its exact locations. Similarly, mapping of Smart-3SEQ reads allowed identification of 3’-end transcript boundaries and polyadenylation sites.

The genome-guided transcriptome assembly SymRos_1_4_RNA.v1 generated by TBONE pipeline contains 34,493 genes, of which 29,351 are protein-coding and 5,142 are non-coding (Table 4). The number of SL trans-spliced genes is 10,433, amounting to 30.2% of all genes. The transcriptome assembly contains 296 out of 303 eukaryotic BUSCO gene models, or 97.6% (Table 4), indicating high quality and completeness of the transcriptome.

**Table 4.**
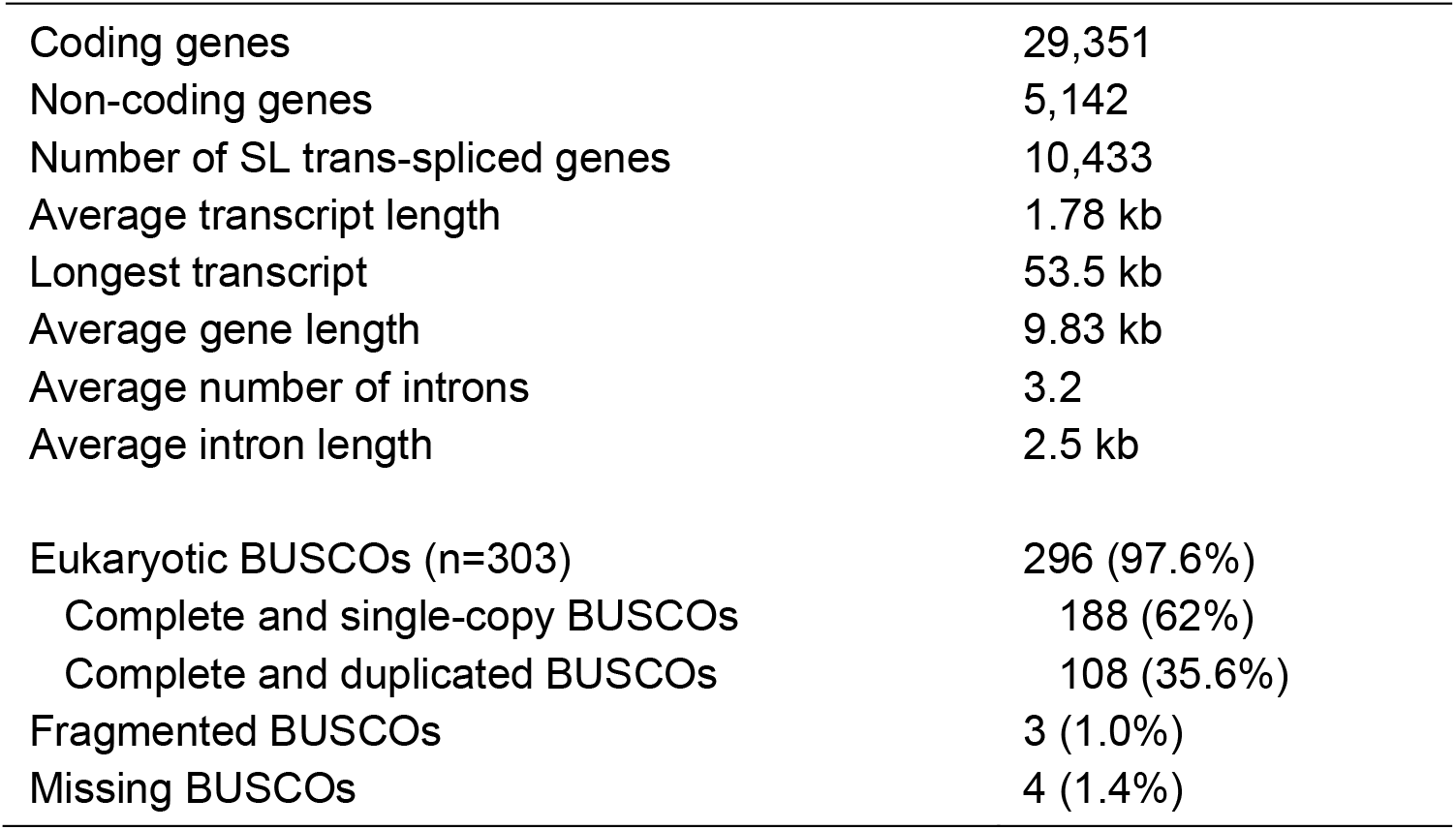
Characteristics of SymRos_1_5_RNA.v1 transcriptome assembly Coding genes 29,351

## Discussion and Conclusions

### General characteristics of the *S. roscoffensis* genome and comparison with other xenacoelomorphs

A draft genome of the acoel *S. roscoffensis* has previously been published based on Illumina sequencing data (Philippe et al. 2019). This was a rather poor-quality genome, assembled in scaffolds with an N50 of less than 5 kb, though it was useful for the initial characterization of small and big gene families (Perea-Atienza et al. 2015; Gavilán et al. 2016). However, the ability of this draft genome to provide useful genomic information was extremely limited. Characters such as indels, exon–intron structures, or syntenic blocks were impossible to identify. The only way to obtain longer scaffolds was to re-sequence the genome using a different method, which in our case was PacBio Hi-Fi technology. Assembly of the Hi-Fi sequence data immediately raised the N50 value by a factor of 40, up to 200 kb, and further genome scaffolding with Hi-C and RNA-seq reads resulted in a highly continuous 1.1 Gb large genome assembly with N50 scaffold size of 1 Mb. Based on flow cytometry we estimated the genome size of *S. roscoffensis* to be in range of 910-940 Mb, which is smaller than the previous estimate of 1.40 Gb (Arboleda et al. 2018), but better corroborates the size of the generated genome assembly. The genome size of *S. roscoffensis* is in the range of those reported for the few sequenced members of this clade: those of *Hofstenia miamia* (950 Mb) (Gehrke et al. 2019) and *Praesagittifera naikaiensis* (654 Mb) (Arimoto et al. 2019). As mentioned above, it is important to note that *S. roscoffensis* lives in symbiosis with the algae *T. convolutae* during its adult stage. The genome described here is only that of the acoel host. An independent, flow cytometry-based analysis of the *T. convolutae* genome has shown that it is markedly smaller than that of the host, by a factor of 7 to 10.

The genome of *S. roscoffensis* is known to be packed into 20 chromosomes of seemingly equal size (2n = 20), as determined cytochemically using chromosomal spreads (Moreno et al. 2009). The current genome assembly has 3,460 scaffolds, with half of the genome represented by 287 scaffolds. While this is a highly continuous genome assembly, it is not at a chromosome level, despite availability of Hi-C data. Failure to generate chromosome-level assembly can be attributed to the highly repetitive nature of the genome, high level of heterozygosity in the population of animals used to generate genome sequencing data and inability of current genome assembly algorithms to efficiently handle such challenging data.

The GC content of this genome amounts to 36.7% (Table 2), so *S. roscoffensis* can be considered as having an AT-rich genetic makeup. The composition fits well in the range observed for other xenacolomorphs (43% GC content in *X. bocki* (Schiffer et al. 2022), or 39.1% GC content in *P. naikaiensis* (Arimoto et al. 2019); all of them, also, enriched in A+T nucleotides.

Gene number has been estimated to be 34,493 genes, with 29,351 of those corresponding to coding and 5,142 non-coding transcripts. The number of identified genes is higher than reported for *H. miamia* (~22,000) or *X. bocki* (~15,000). This variability can be explained by differences in annotation pipelines used to identify genes, with TBONE pipeline used in this study being more inclusive for non-conserved, non-coding, repetitive and low-expressed genes.

Many organisms have shown to process transcripts by trans-splicing, a process that connects exons of two different primary RNA molecules transcribed from *a priori* unrelated genomic loci (Lasda and Blumenthal 2011). In several of those organisms, a dominant form of trans-splicing is SL trans-splicing, where the same spliced leader (SL) sequence is trans-spliced to different mRNAs (Lasda and Blumenthal 2011). Interestingly, we have observed that 30.2% of the genes in *S. roscoffensis* are SL trans-spliced (Table 3), which is similar to the number of trans-spliced genes observed in the flatworm *Macrostomum lignano* (Ustyantsev and Berezikov 2021). The prevalence of trans-splicing events ranges from 1.58% in groups such as insects (Kong et al. 2015) to ~80% in the ascidian *Ciona intestinalis* (Matsumoto et al. 2010). This wide range can also be explained, in part, by the use of different analytical tools, not capturing the real frequency with which different transcripts participate in trans-splicing events (Matsumoto et al. 2010). Further analysis has to be carried out in order to clarify which, when and how frequently, the *S. roscoffensis* genes participate in SL trans-splicing events.

### Repeated elements in the *S. roscoffensis* genome and commonalities with other xenacoelomorph genomes

Large fractions of eukaryotic genomes consist of repetitive elements, which vary considerably in their abundance across species. The repetitive fraction of the genome, known as the *repeatome*, correlates with genome size both within and among species (Lynch 2007) and therefore likely plays a major role in genome size evolution (Talla et al. 2017). In the assembled genomes of acoels, the content of repetitive sequences varies from a value of 53% in *H. miamia* to 61% in *S. roscoffensis* and the largest estimated value of 70% in *P. naikaiensis*. Moreover, in the related group of xenoturbellids, a recent sequencing project has revealed that the species *X. bocki* has a surprisingly low fraction of repeats, around 25% (Schiffer et al. 2022). Interestingly, *X. bocki* also has a relatively compact genome of ~110 Mb (Schiffer et al. 2022). In fact, the fraction of repeats is directly correlated with the estimated size of the genomes and in cases of *S. roscoffensis* and *H. miamia* it is in the same range, though *P. naikaiensis* is an example that clearly deviates from those values and ratios. In this last case the perhaps anomalous reported value might be the result of a genome assembled with shorter scaffolds and, thus, in need of further refinement.

In all acoel genomes a big fraction of the repetitive sequences are retrotransposons (41.5% in *P. naikaiensis* or 51.75% in *S. roscoffensis*), and some of them have already been localized around, for instance, the Hox transcriptional units (and probably responsible of the disintegration of the acoel HOX cluster; see (Moreno et al. 2011)). Our current knowledge is, nowadays, quite limited, and it is clear that further studies are needed to improve our understanding of the diversity of retrotransposons across the Xenacoelomorpha, plus the evolutionary timing of their respective insertions.

## Materials and Methods

### Preparation of aposymbiotic animals

Aposymbiotic animals were used as a source of genomic DNA in order to avoid sequences of microalgae symbionts. For this, gravid animals were collected at low tide from beaches in the areas of Roscoff and Carantec, Brittany, France and were transported to laboratory, where most of them spontaneously spawned. For samples from Carantec, the resulting cocoons were reared in an environmental chamber at 15°C with a 12h–12h light–dark cycle, and hatched juveniles were stored at −20°C in RNAlater until DNA and RNA extractions could be performed. For the Roscoff samples, the resulting cocoons were reared in an environmental chamber at 13°C without light as an additional measure to prevent microalgae growth, and juveniles were snap-frozen in liquid nitrogen and transported on dry ice for chromatin extraction.

### Genome size measurement

The genome size of *S. roscoffensis* was determined by a flow cytometry approach (Hare and Johnston 2011) as previously described for *Macrostomum lignano* flatworms (Wudarski et al. 2017). Worms were collected in an Eppendorf tube, medium was aspirated, and 200 µl 1x Accutase (Sigma, A6964-100ML) was added. The samples were incubated at room temperature for 30 min, followed by tissue homogenization through pipetting to obtain a cell suspension. 800 µl f/2 medium was added to the suspension and cells were pelleted by centrifugation at 4°C, 1000 rpm for 5 min. The supernatant was then aspirated and the cell pellet was resuspended in the nuclei isolation buffer (100 mM Tris-HCL pH 7.4, 154 mM NaCl, 1 mM CaCl_2_, 0.5 mM MgCl2, 0.2% BSA, 0.1% NP-40 in MilliQ water), followed by more pipetting to obtain a nuclei suspension. This treatment was performed on samples with adults, which include symbionts, and juveniles without symbionts. *Macrostomum lignano* NL12 line worms (Wudarski et al. 2017), which served as a positive control, were treated in the same way. Human fibroblasts, which served as reference, were treated in the same way as the cell suspensions of the worms. All nuclei suspensions were passed through a 35 µm pore size filter (Corning, 352235) and treated with RNase A and 10 µg/ml PI for 15 min prior to measuring PI fluorescence with a BD FacsCanto II Cell Analyzer. All samples were first measured separately, and then as combination of samples.

### Pacific Biosciences Hi-Fi genome sequencing

Genomic DNA from aposymbiotic animals stored in RNAlater was extracted using MagAttract High Molecular Weight DNA extraction kit from Qiagen following manufacture’s protocol with some modifications: lysis of the sample was performed at 37°C overnight without shaking, and subsequent washing steps were performed at 800 rpm.

The preparation and sequencing of the genomic library was performed by GenomeScan B.V. (Leiden, The Netherlands). To begin, gDNA (15 μg) was fragmented to the desired length of ~15 kb using Covaris g-Tubes. The concentration of the fragmented DNA was determined using a Qubit2.0 fluorometer. The sample was then diluted to 0.3 ng/μl for quality control using the genomic DNA 165 kb kit on the Femto Pulse system. HiFi SMRTbell libraries were developed using PacBio’s SMRTbell Express Template Prep Kit 2.0 protocol using 10 μg of fragmented DNA input. The fragmented DNA was end-repaired, A-tailed, and ligated with a PacBio hairpin adapter to create a SMRTbell library. Nuclease treatment of the library was performed to remove damaged or non-intact SMRTbell templates. Using the same procedure as for the initial fragmented DNA, the concentration of the SMRTbell library was determined using a fluorometer and the sample was then diluted to 0.3 ng/μl for quality control. BluePippin was used to select two size fractions, 9–13 kb and >15 kb, from the SMRTbell library for HiFi long-read sequencing. Based on the Fragment Analyzer (Femto Pulse) results, the fraction with fragments >15 kb was selected for further HiFi long-read sequencing. Sequencing of the sample was performed using diffusion loading on the Sequel II SMRT Cell 8M with Sequel II Sequencing Kit 2.0 reagents. The SMRTcell (SMRT Cell 8M) was sequenced for 30 hours. The obtained raw reads were processed with the ccs tool v.6.2.0 (RRID:SCR_021174) to generate circular consensus sequences, and further filtered for remaining adapter sequences using HiFiAdapterFilt v. 2.0.0 (Sim et al. 2022).

### Hi-C library construction and sequencing

The preparation and sequencing of Hi-C library was performed by Arima Genomics (San Diego, USA) at the Arima High Coverage Hi-C Service. For this, aposymbiotic animals from a dense culture were concentrated in the volume of 150 μl with as much medium removed as possible, snap-frozen and shipped on dry ice to Arima Genomics. The library was prepared using the Arima-HiC+ kit and the Arima Library Prep Module according to the manufacturer’s protocols. This library was, then, sequenced on a Illumina HiSeq X instrument.

### RNA library construction and sequencing

Aposymbiotic animals were collected and stored at −20°C in RNAlater until RNA isolation. RNA was isolated with Qiagen RNeasy Micro Kit according to the manufacturer’s protocol, except that the DNase I treatment step was omitted. Preparation of the RNA library was performed according to Smart-3SEQ protocol (Foley et al. 2019). The library was sequenced on the Illumina NextSeq 500 instrument.

### *De novo* transcriptome assembly

*De novo* transcriptome assembly was generated using ReCAP assembly pipeline (Grudniewska et al. 2016). Publicly available *S. roscoffensis* RNA-seq data (SRR5760179 and SRR8506641) were normalized to 30x coverage and assembled into initial contigs using Trinity v.2.11.0 package (Grabherr et al. 2011) with default parameters and accounting for strand information. RNA-seq reads were mapped to the contigs using Bowtie v.2.3.4.3 (Langmead and Salzberg 2012), and reads mapping to a given contig extracted and reassembled using CAP3 v 12/21/07 (Huang and Madan 1999) requiring consistent placing of paired-end reads. The redundancy of the resulting transcriptome was reduced by prioritizing transcripts by read mapping and selecting the minimal transcript set that explains all mapped RNA-seq reads. The resulting *de novo* transcriptome assembly was named SYMROS200831.

### Genome assembly and evaluation

PacBio Hi-Fi data were assembled independently with FALCON/FALCON-Unzip v.1.8.1/v.1.3.7 (Chin et al. 2016), Flye 2.9 (Kolmogorov et al. 2019), HiCanu v.2.2 (Nurk et al. 2020), Hifiasm v.0.16.1 (Cheng et al. 2021), IPA v.1.8.0 (Sovic 2022), Peregrine v.0.1.5.3 (Chin and Asif Khalak 2019), Raven v.1.8.1 (Vaser and Šikić 2021) and wtdbg2 v.2.5 (Ruan and Li 2020) using parameters default for each assembler. Assemblies with the resulting size greater than the measured genome size were de-duplicated by purge_dups v.1.2.5 (Guan et al. 2020). The quality of the assemblies was evaluated by mapping *de novo* transcriptome assembly SYMROS200831 to the genome assemblies with GMAP v.2021-08-25 (Wu and Watanabe 2005) and calculating the fraction of transcripts that map to a given assembly with at least 90% of transcript length covered and the fraction of eukaryotic BUSCO models (v.2.0) present (Simão et al. 2015).

Hi-C reads were mapped to the deduplicated peregrine assembly by BWA aligner v.0.7.17-r1188 (Li and Durbin 2009), and processed by Arima Genomics mapping pipeline (https://github.com/ArimaGenomics/mapping_pipeline). Hi-C genome scaffolding was performed by SALSA2 v.2.3 software (Ghurye et al. 2019). Paired-end RNA-seq reads (SRR5760179 and SRR8506641) were mapped to the scaffolds with HISAT v.2.2.1 (Kim et al. 2019) and further scaffolding was performed by P_RNA_scaffolder (Zhu et al. 2018). Gap closing was performed with LR_gapcloser v.1.1 (Xu et al. 2019) using initial PacBio Hi-Fi reads. Final polishing was done by pilon v.1.24 (Walker et al. 2014) with RNA-seq reads. The resulting genome assembly was named SymRos_1_5.

### Mitochondrial genome assembly and annotation

The mitochondrial genome was reconstructed by performing tblastn (Camacho et al. 2009) searches in genome assemblies generated by different assemblers using 11 protein coding sequences from the *Isodiametra pulchra* mitochondrial genome (NCBI Reference Sequence NC_034948.1). The maximum number of blast hits (8/11) was found in the FALCON assembly, and two contigs (99.5% identical) of length 14832 bp and 15353 bp were extracted. Contigs were analysed by self-blast, a single 100% identical terminal repeat (length 535) corresponding to COXI gene was identified in one of the contigs, and the contig was circularized using this repeat information. The resulting mitochondrial genome was annotated using MITOS2 web-server (Donath et al. 2019) and compared to the previously published (Mwinyi et al. 2010) *S. roscoffensis* mitochondrial genome (NCBI Reference Sequence NC_014578.1) using BLAST (Camacho et al. 2009).

### Repeat analysis

Tandem repeats were annotated and masked with Tandem Repeat Finder (TRF) 4.10.0 (Benson 1999). A *de novo* library of classified repetitive element models was created using RepeatModeler 2.0.3 (Flynn et al. 2020). Homology-based annotation of full-length and fragmented long terminal repeat (LTR) retrotransposons was done using the Domain-Associated Retrotransposon Search (DARTS) algorithm (Biryukov and Ustyantsev 2021). Each full-length or partial sequence identified by DARTS was added to the custom LTR retrotransposon library. RepeatModeler- and LTR retrotransposon-derived libraries were merged together and used as input for RepeatMasker 4.1.2-p1 (RepeatMasker, RRID:SCR_012954) (Tempel 2012) to map locations of classified repetitive sequences in the genome.

### Gene prediction and annotation

Gene annotation was performed using TBONE pipeline (Wudarski et al. 2017), which facilitates identification of trans-spliced genes. Publicly available RNA-seq data SRR5760179 and SRR8506641 and Smart-3SEQ data generated in this study were mapped to the genome assembly with HISAT v.2.2.1 (Kim et al. 2019). Resulting bam files were used to generate initial gene models independently by Scallop v0.10.5 (Shao and Kingsford 2017) and StringTie v2.2.1 (Kovaka et al. 2019). *De novo* transcriptome assembly SYMROS200831 was mapped to the genome assembly with GMAP v.2021-08-25 (Wu and Watanabe 2005). GTF files resulting from Scallop and StringTie assemblies and from the *de novo* transcriptome mapping were merged by gffread v.0.12.7 (Pertea and Pertea 2020). The resulting transcriptional units were further processed to identify trans-spliced genes.

Trans-splicing leader sequence was determined through the analysis and mapping of the *de novo* transcriptome assembly SYMROS200831 to the genome assembly. Transcripts, in which first 10-50 nucleotides are not mapped to the genome were identified and the respective non-mapped 5’ sequences were extracted from the transcripts and sorted by the number of transcripts in which they were present. The most abundant sequences were manually aligned to each other and the trans-splicing leader sequence GCCTAATTGTTGTGATAAACTTATTAAATAGA was reconstructed from this alignment. To determine the SL RNA gene structure in the genome assembly, we used this sequence as a seed for a blastn (Camacho et al. 2009) search requiring 100% matching hits. A conserved overlapping region between all the BLAST hits of ~100 bp was extracted and examined for canonical SL RNA folding using the RNAfold web server (Gruber et al. 2008).

Reads containing trans-splicing sequence were extracted form RNA-seq data (SRR5760179 and SRR8506641), trimmed and mapped to the genome assembly with HISAT v.2.2.1 (Kim et al. 2019). The resulting wiggle files were used to identify genomic peaks corresponding to trans-splicing locations. Similarly, peaks corresponding to polyadenylation sites at 3’ end gene boundaries were identified by mapping reads from Smart-3SEQ RNA-seq libraries, which capture 3’ ends of transcripts. The generated trans-splicing and polyadenylation signals were overlapped with the genomic coordinates of transcriptional units to refine gene boundaries and separate trans-spliced genes. Open reading frames (ORFs) were predicted by TransDecoder v.5.5.0 (Haas et al. 2013) and 3’UTRs extended by merging adjacent transcripts in cases where ORF-containing transcripts was missing poly(A) signal but downstream transcript had poly(A) signal but did not have predicted ORF, and the transcripts were separated by a region of not more than 2 kb, 5 kb or 10 kb containing at least 0%, 30% or 90% of repetitive sequences respectively. The resulting transcriptome annotation was named SymRos_RNA_1_5.v1 and includes alternative splicing forms as well as non-coding transcripts. To remove redundancy, for each genomic locus a single representative transcript was selected and included into a subset called ‘core genes’, which is used in the majority of downstream analyses.

## Supplementary Material

**Supplementary Figure 1:**
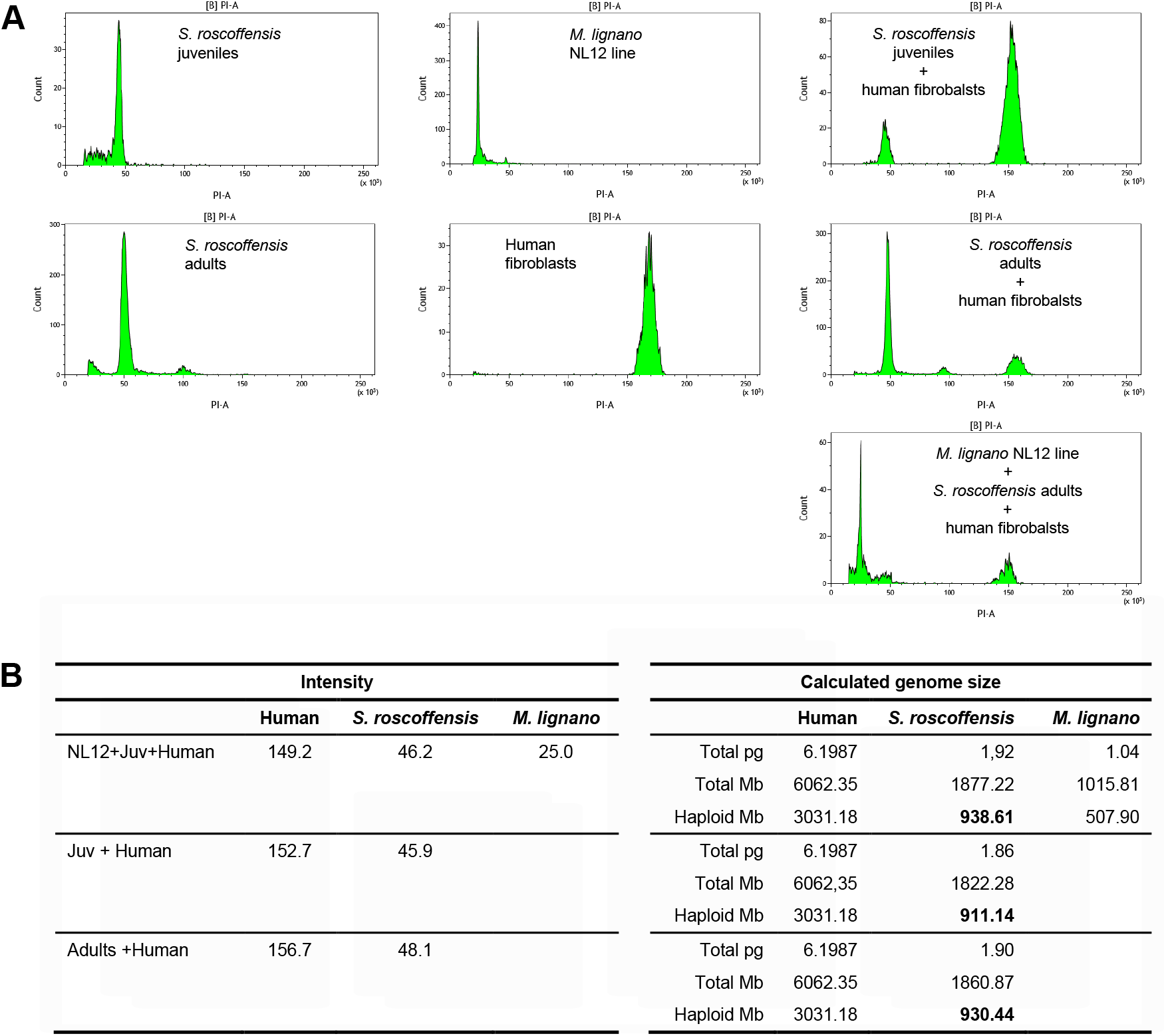
Genome size measurement of *S. roscoffensis*. (A) Separate and combined measurements of fluorescence in *S. roscoffensis, M. lignano* NL12 line and human fibroblasts. (B) Calculation of *S. roscoffensis* genome sizes using human fibroblasts as a reference and *M. lignano* NL12 line as a positive control.

**Supplementary Figure 2:**
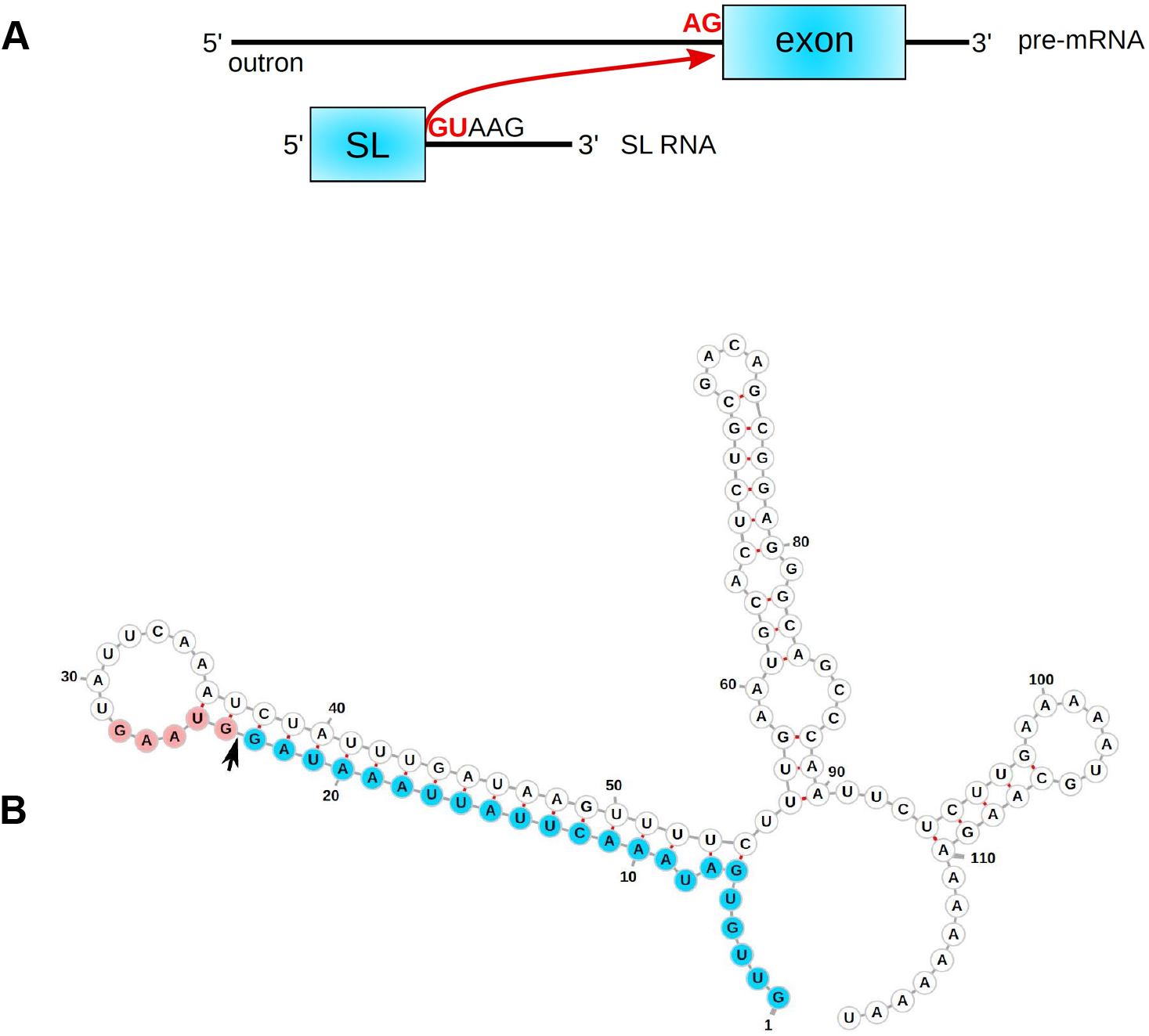
SL trans-splicing in *S. roscoffensis*. (A) General principle of SL trans-splicing. Spliced leader (SL) sequence from SL RNA is trans-spliced to an exon of a target pre-mRNA. (B) Predicted secondary structure of putative *S. roscoffensis* SL RNA. Spliced leader sequence is show in blue. Arrow indicates the site of trans-splicing.

## Acknowledgements

We would like to acknowledge Brenda Gavilán and Sergio Melero for helping us with the collection of hatchlings. The work in P. Martinez’s laboratory was funded by the Ministerio de Ciencia, Innovación y Universidades, Spain (project number: PGC2018-094173-B-I00). The work in E. Berezikov’s laboratory was supported by the Dutch Research Council Open Competition XS grant (file number OCENW.XS21.2.051). The work of M. Biryukov on repeat annotation was supported by the Russian State Budget project No. 0259-2021-0009. S. G. Sprecher was supported by Swiss National Science Foundation grant 310030_188471 and IZCOZ0_182957. X. Bailly was funded by the Functional Genomics Joint Research Activities of the ASSEMBLE Plus Program (Association of European Marine Biological Laboratories Expanded).

## Author contributions

PM, XB, SGS and EB conceived the project. PM and EB designed and supervised the project. PM, SGS and XB provided materials. LG and SM isolated genomic DNA, made RNA-seq libraries and measured genome size. EB generated genome and transcriptome assemblies. KU and MB annotated repeats and trans-splicing. PM and EB provided the funds and wrote the paper. All authors read and revised the final version of the manuscript.

## Data Availability

All raw sequencing data have been deposited in the NCBI Sequence Read Archive (accession codes SRR20990873 - SRR20990875) and can be accessed with BioProject No. PRJNA867535. The genome assembly has been deposited at DDBJ/ENA/GenBank under the accession JANVAR000000000. The version described in this preprint is version JANVAR010000000. The annotated genome is available via UCSC Genome Browser interface at http://gb.macgenome.org.

## Notes

### Competing Interest Statement

The authors have declared no competing interest.

